# Versatile toolkit for highly-efficient and scarless overexpression of circular RNAs

**DOI:** 10.1101/2023.11.21.568171

**Authors:** Brett W. Stringer, Marta Gabryelska, Shashikanth Marri, Letitia Clark, He Lin, Laura Gantley, Ryan Liu, Jeremy E. Wilusz, Vanessa M. Conn, Simon J. Conn

## Abstract

Circular RNAs (circRNAs) are a class of single-stranded, covalently closed RNA that contain a unique back-splice junction (bsj) sequence created by the ligation of their 5’ and 3’ ends via spliceosome-catalyzed back-splicing. A key step in illuminating the cellular roles of specific circRNAs is via increasing their expression. This is frequently done by transfecting cells with plasmid DNA containing cloned exons from which the circRNA is transcribed, flanked by sequences that promote back-splicing. We observed that commonly used plasmids lead to the production of circRNAs with molecular scars at the circRNA bsj. Stepwise redesign of the cloning vector corrected this problem, ensuring *bona fide* circRNAs are produced with their natural bsj at high efficiency. The fidelity of circRNAs produced from this new construct was validated by RNA sequencing and also functionally validated. To increase the utility of this modified resource for expressing circRNA, we developed an expanded set of vectors incorporating this design that (i) enables selection with a variety of antibiotics and fluorescent proteins, (ii) employs a range of promoters varying in promoter strength and (iii) generated a complementary set of lentiviral plasmids for difficult-to-transfect cells. These resources provide a novel and versatile toolkit for high-efficiency and scarless overexpression of circular RNAs that fulfill a critical need for the investigation of circRNA function.

## INTRODUCTION

Circular RNA (circRNA), as the name implies, is RNA that exists in a covalently-closed state rather than the canonical, linear form (1). This property enhances its stability and increases its half-life compared to linear RNA (2), yet despite this, circRNA is a rare form of RNA, accounting for just 0.1% of total RNA within eukaryotic cells (3). The vast majority of circRNAs are formed by a process known as back-splicing, in which the splicing machinery joins the 3’ end of an exon to the 5’ end of the same or an upstream exon, thereby forming a closed circle of single-stranded RNA. This process creates a sequence at the joined ends, known as the back-splice junction (bsj), that is not present in the linear RNA molecule. CircRNAs formed in this manner are a theoretical possibility for any gene with multiple exons, yet not all transcribed, spliced genes produce circRNAs (4). Indeed, the expression of circRNA is dynamic and tightly regulated, with some conserved across species, and their expression can be independent of the parental mRNA (5–7). This ancient and conserved gene expression phenomena implies functions for circRNAs.

The structure of RNA has been previously shown to be critical for its functional interactions with other RNAs and protein targets within the cell. This includes functional folding of catalytic RNAs (eg. ribozymes) (8), interactions of viral RNAs with host cellular components (9) and the ability of microRNAs to bind the 3’ untranslated region of messenger RNA (10). One of the special features of RNA is flexibility, making its structure responsive and strongly affected by the cellular environment (11, 12). The overall flexibility of RNA strands directly reflects the universal rule of sequence-structure-function relationship. Importantly, Liu *et al*. demonstrated that circular RNA and linear RNA comprising the same sequence can show distinct structures, with circRNA intrinsic RNA duplexes playing functional roles in innate immunity (13). Therefore, it is of utmost importance to assure fidelity of the sequence of any overexpressed RNA, including circRNA.

The circRNA bsj is a unique series of nucleotides which is known to affect functional interactions with RNA, DNA or protein targets, alter RNA conformation, and/or influence the open reading frames of protein-coding circRNAs. One report found that *circTTN1* contains a sequence motif spanning the bsj that binds SRSF10, impacting the ability of SRSF10 to regulate splicing (14). Loss of this circRNA resulted in abnormal splicing of important cardiomyocyte SRSF10 targets such as *MEF2A* and *CASQ2*, and abnormal splicing of *TTN* itself. This resulted in structural abnormalities and apoptosis in human induced pluripotent stem cell-derived cardiomyocytes and reduced contractile force in engineered heart tissue. Another notable report demonstrated that *circZNF609* can be translated as it encodes an open reading frame which spans the bsj, with functional role in myogenesis (15), and a number of other circRNAs have been proposed to contain ORFs that similarly span the bsj (16). These examples highlight how circRNA function can be specifically affected by the bsj, demanding a high-fidelity system for overexpression to maintain functional relevance.

Due to its low abundance, investigation of circRNA function commonly includes ectopically overexpressing the circRNA (17). There are now several molecular biology strategies to achieve circularization of RNA *in vitro* and/or *in vivo* including the use of cognate self-complementary introns, self-splicing introns and ribozymes (18). The inaugural approach for generating circular RNA overexpression constructs was to replicate the native genomic DNA context, with cognate introns and exons cloned into a single construct (19). This native milieu was modified to augment circularisation efficiency by including a copy of the 5’ intron region (typically 100-750 nucleotides in length) downstream of the circRNA exon in a reverse complement orientation, thereby producing a long hairpin structure between the introns that promotes circularisation of the intervening exons. This approach has yielded high fidelity circRNA overexpression in most instances, including our own laboratory, but this strategy demands multiple PCR, restriction digestion, ligation and sequencing verification steps (7, 20, 21). Furthermore, due to the long reverse complementary stretch of the amplicon, it is not amenable to high-throughput cloning strategies, including use of synthetic gene fragments.

In order to address these concerns, a number of plasmid-based “universal” circRNA overexpression constructs have been published (22–24). This strategy involves cloning exons which comprise the circRNA into a multiple cloning site flanked by partial intron sequences containing reverse complementary regions. This approach is designed to simplify cloning while facilitating splicing and promoting back-splicing. The most widely utilised construct, the pcDNA3.1(+) CircRNA Mini Vector (Addgene plasmid #60648) (Figures 1A-B), hereafter referred to as “mc”, incorporates portions of native introns from the human *ZKSCAN1* gene that contain inverted *Alu* repeat elements (22). Liang and Wilusz (22) demonstrated that this vector successfully circularised *circZKSCAN1(2,3)* by Sanger sequencing of RT-PCR products and Northern blotting. While this was an anticipated result as it is the native introns driving circularisation of its cognate circRNA, this report also demonstrated that other circRNA products, e.g. from the human *EPHB4* gene, could be generated from the construct (22). The mc plasmid has been further used in numerous reports since to achieve high levels of overexpression of non-*ZKSCAN1* circRNAs. However, 15 out of 17 systematically checked citations since 2014 did not report the bsj sequence of the overexpressed circRNA, either by Sanger sequencing of RT-PCR products or high-throughput RNA sequencing to verify the scarless bsj, despite it being considered best-practice for circRNA research (25). As a result, there is a possibility that conclusions in these reports may be confounded. While circRNA function may not be solely confined to the bsj, investigations that have studied circRNAs containing vector sequence may have failed to identify a bsj-related activity, incorrectly ascribed a function to an otherwise inert circRNA or incorrectly concluded a circRNA lacked function, as a result of the integrity of the bsj being compromised.

**Figure 1.**
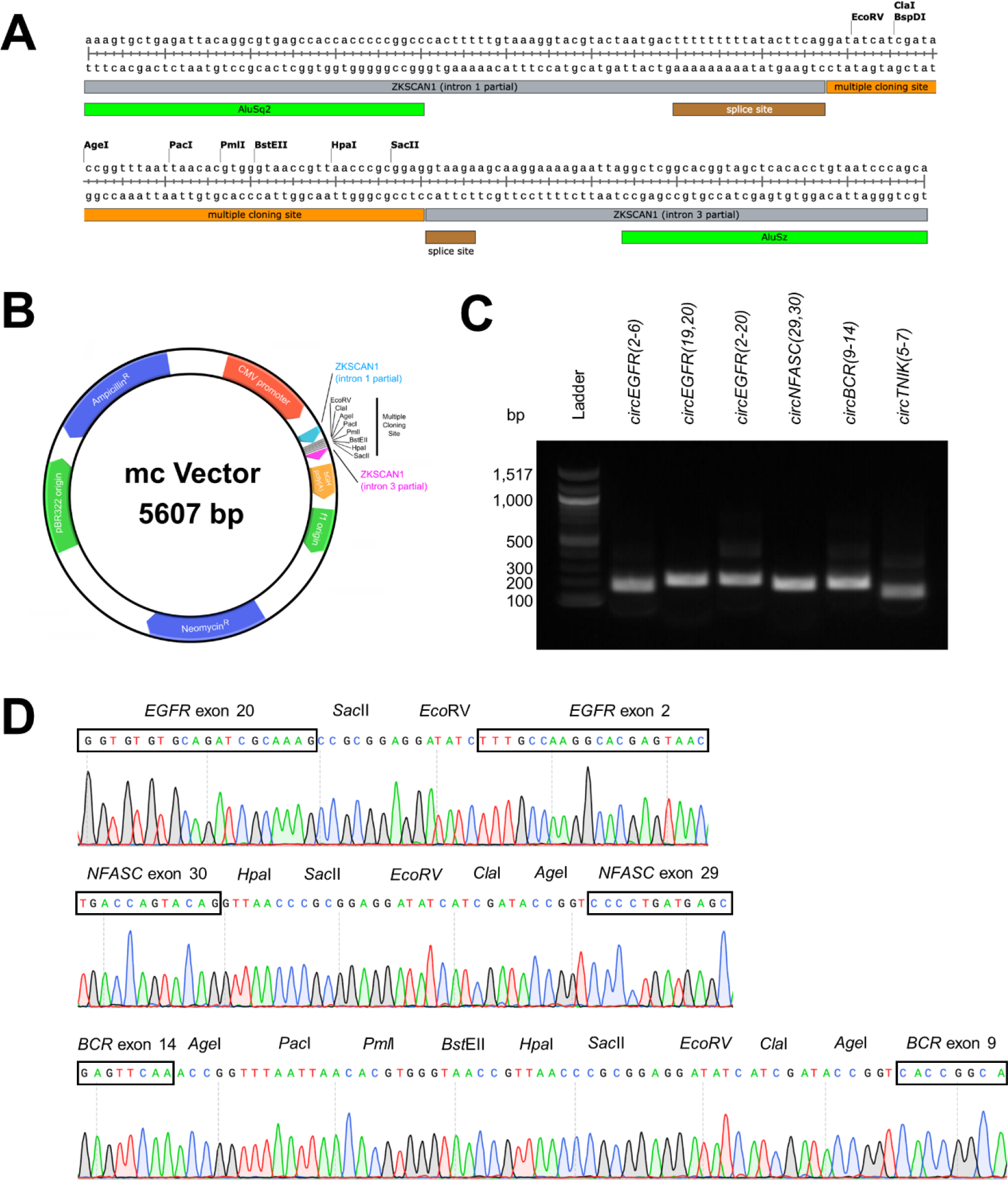
CircRNA Mini Vector incorporates vector multiple cloning site sequence within the back-splice junction of expressed circRNAs. (**A**) Nucleotide sequence of the pcDNA3.1(+) CircRNA Mini Vector (called mc) multiple cloning site and flanking intronic regions used to produce circRNA from cloned exons. (**B**) Plasmid map of CircRNA Mini (mc) vector generated using PlasMapper 3.0 (34). (**C**) Back-splice junction RT-PCR amplicons from HEK293 cells transiently transfected with mc vector clones of *circEGFR(2-6), circEGFR(19,20), circEGFR(2-20), circNFASC(29,30), circBCR(9-14)* and *circTNIK(5-7)* visualised by agarose gel electrophoresis. (**D**) Sanger sequencing chromatograms of circRNA bsj sequence of RT-PCR amplicons from HEK293 cells transfected with mc vector clones of *circEGFR(2-20), circNFASC(29,30)* and *circBCR(9-14)*. Restriction sites encompassing the multiple cloning site of mc vector written above sequence, with circRNA bsj sequence in boxes.

Through our extensive investigation of a panel of circRNAs cloned into this plasmid it was found to incorporate unwanted vector sequence into the bsj of each circRNA transcript. Consequently, we sought to correct this issue by re-engineering the plasmid, thereby producing circRNAs with *bona fide* bsj with high-efficiency and fidelity. Herein we report the construction and validation of the redesigned vector, named mc2, and an expanded set of plasmids derived from it to facilitate the investigation of circRNA function, including the role of the unique bsj.

## MATERIAL AND METHODS

### Plasmid construction

**“**CircRNA Mini” expression plasmids (herein also abbreviated to “mc”) were derived from pcDNA3.1(+) CircRNA Mini Vector (Addgene plasmid #60648; RRID:Addgene_60648). Exon sequences, amplified by PCR from cDNA or gBlocks synthetic gene fragments (Integrated DNA Technologies), were cloned between the *Eco*RV and *Sac*II, *Age*I and *Hpa*I, or *Cla*I and *Pac*I restriction sites pairs of CircRNA Mini Vector by restriction digestion (New England Biolabs) and DNA ligation using T4 DNA Ligase (New England Biolabs).

To generate the mc2 vector, the multiple cloning site of CircRNA Mini Vector was replaced with a pair of inverted *Bsm*BI restriction sites, oriented such that *Bsm*BI restriction digestion completely removes the intervening vector sequence between the flanking intronic sequences that promote circularisation. Nucleotide sequences to be transcribed as RNA and subsequently circularised can be cloned by one of four methods (Supplementary Figure S2). Plasmid DNA was isolated from overnight cultures (37°C, 230 rpm, 20 hours) of chemically competent E. coli (TOP10- or Stbl3) using Isolate II plasmid miniprep kit (Bioline).

Plasmid sequences were confirmed by Sanger sequencing with the BigDye™ Terminator v3.1 Cycle Sequencing Kit (ThermoFisher Scientific), purified by the BigDye XTerminator™ Purification Kit (ThermoFisher Scientific), separated by SeqStudio™ Cartridge v2 (ThermoFisher Scientific) on the SeqStudio Genetic Analyzer (ThermoFisher Scientific) and/or long read sequencing on the MiniON Mk1C (Oxford Nanopore) using the Rapid Barcoding Kit (Oxford Nanopore) on the Spot On Flow Cell Mk1 R9 Version (Oxford Nanopore). List of constructs used are shown in Supplementary Table S2.

### Cell culture

Cell lines (HEK293, HeLa and HepG2) were cultured as adherent monolayers in filter-capped plastic flasks (Corning or Sarstedt) in DMEM (Thermo Fisher Scientific) supplemented with 10% foetal bovine serum (Bovogen) and 1× Antibiotic-Antimycotic (Gibco). Cells were cultured in 5% CO_2_/95% humidified air at 37°C. Cell identity was verified by short tandem repeat profiling (AGRF, Melbourne) and all cell lines were routinely tested for mycoplasma and confirmed to be mycoplasma-free.

### Cell transfection

Cell lines were transfected at 70-80% confluence in 12-well plates (Corning or Sarstedt) using Lipofectamine™ 2000 transfection reagent (Thermo Fisher Scientific) as recommended by the manufacturer (4 μL Lipofectamine™ 2000 and 1.6 μg plasmid were each added to 100 μL Opti-MEM and combined per transfection). 1.5 × 10^5^ cells were plated per well in 1 mL DMEM plus 10% FBS 24 hours prior to transfection.

### Cell transduction

Cell lines in 6-well tissue culture plates (Corning or Sarstedt) in 3 mL of medium were transduced when 70-80% confluent with 1 mL of unconcentrated lentivirus followed by incubation at 37°C. Virus-containing medium was replaced next day with fresh medium and antibiotic selection commenced 48 hours after lentiviral transduction began.

### Quantitative real-time PCR

Total RNA was isolated from cells lysed *in situ* using TRIzol (Invitrogen) and purified with a Direct-zol RNA miniprep kit (Zymo Research), using on-column DNaseI digestion of genomic DNA. cDNA was prepared from 1 μg of total RNA using a QuantiTect™ Reverse Transcription kit (Qiagen) from random hexamer primers. Real time quantitative PCR was performed by Quantitect SYBR Green PCR kit (Qiagen) with a RotorGene™ Q instrument (Qiagen). Data were analysed by relative quantitation using the comparative C_T_ method and normalised to *TBP*. PCR primers with melting temperatures of approximately 60°C were designed using Primer3 to amplify unique PCR products 100-300 bp long. Gene-specific oligonucleotide sequences are shown in Supplementary Table 1. Q-PCR products were visualised by agarose gel electrophoresis, recovered using a QIAquick gel extraction kit (Qiagen) and sequenced by Sanger sequencing to confirm their identity and to determine the sequence of the back-splice junction.

### DNA:RNA Immunoprecipitation sequencing (DRIP-seq)

DRIP-seq was performed on HEK293 cells as per Conn *et al*. (21) compared to input DNA.

### CircRNA back-splice junction amplicon-seq

HEK293 cells in 12-well plates were transfected with 1.6 μg of CircRNA Mini or mc2 plasmid and total RNA was isolated 24 hours later as described above in 35 μL of water. Twenty microlitres of total RNA was digested with 2.5 units of RNase R (Biosearch Technologies, RNR07250) for one hour at 37°C to remove linear RNA. CircRNA was purified using RNeasy MinElute columns (Qiagen) as recommended by the manufacturer and eluted in 14 μL of water. Reverse transcription was performed with 12 μL of circRNA as described above, using circRNA-specific primers appended with 12-nucleotide unique molecular identifier (UMI) and Illumina reverse adapter sequences (see Supplementary Table 1 for oligonucleotide sequences). A single round of PCR was then performed using Q5 high fidelity DNA polymerase (NEB) with exon-specific primers appended with Illumina forward adapter sequence. Exonuclease digestion was then performed with thermolabile exonuclease I (NEB, M05685) to remove excess primers (37°C for 4 minutes then 80°C for 1 minute). Twenty-five cycles of PCR were then performed using Q5 high fidelity DNA polymerase (NEB) and Illumina forward and reverse adapter primers (98°C for 10 s, 60°C for 20 s, 72°C for 10 s). The reactions were then subjected to agarose gel electrophoresis (1% agarose, 1× TAE, 20 V/cm, 30 mins). Amplicons (∼250 bp) were excised from the gel, recovered using Qiaquick (Qiagen), and eluted in 30 μL of water (7-14 ng/μL final concentration). Libraries and sequencing were prepared by the South Australian Genomics Centre (SAGC) using the Illumina 16S Metagenomic Sequencing Library Preparation Protocol with the following changes: 10 μL of Amplicon product was purified using a 1:0.8 ratio of XP beads (Beckman Coulter), with beads resuspended in a final volume of 10 μL of nuclease-free water. For indexing, 10 μL of cleaned amplicon product was used with 5 μL of IDT UDI indexes (Integrated DNA Technologies) per sample. Final products were quantified by Nanodrop nanospectrophometer (ThermoFisher Scientific) and separated by LabChip (Perkin Elmer) using the HT DNA NGS 3K Reagent Kit (Revvity) to confirm library size and quantities. Libraries were pooled at equimolar concentrations and run on the MiSeq Micro V2 300 cycle kit (Illumina, PE150) using 5 pM with 20% PhiX. Unique sequence reads from individual circRNAs were identified using UMI sequences and back-splice junctions were investigated for the presence of scar sequences.

### Luciferase assay for circRNA functional assessment

Scarred or clean bsj of *circMLLT2(3,4)* was cloned into the SV40 early promoter of psiCheck2 (Promega) by digesting the plasmid with *Stu*I and *Nhe*I (New England Biolabs) using gBlocks synthetic DNA fragments (Integrated DNA Technologies) and NEBuilder HiFi DNA assembly mix (New England Biolabs). HEK293 cells in 12-well plates were transfected with 1.6 μg of CircRNA Mini or mc2 plasmid encoding *circMLLT2(3,4)* and co-transfected with scarred bsj or clean bsj cloned into psiCheck2, respectively. Luciferase assay was performed using the Dual-Glo® Luciferase Assay System (Promega) in triplicate and assay activity was measured with luminescence with the ClarioStar plus spectrophotometer (BMG Labtech).

### Statistics

All statistical analyses were performed using GraphPad Prism 10. Standard error of the mean was reported unless otherwise stated. Significance, where determined, was assessed as described in the Figure legends. Significance level (alpha) was determined as P < 0.05.

## RESULTS

### CircRNA Mini Vector incorporates molecular scars into the circRNA back-splice junction

To assess the CircRNA mini vector, hereafter called mc, six circRNAs were restriction enzyme-cloned into this vector – *Eco*RV/*Sac*II cloning sites: *circEGFR(2-20), circEGFR(2-6), circEGFR(19-20), circTNIK(5-7)*; *Hpa*I/*Age*I cloning sites: *circNFASC(29-30)*, and using the *Age*I site only: *circBCR(9-14)* (Figures 1A-B). All plasmid sequences were confirmed by Sanger sequencing. These plasmids were transfected into HEK293 cells and 24 hours post transfection, RNA was isolated and circRNA expression was assessed by RT-PCR and gel electrophoresis. Each construct yielded a single amplicon of the anticipated size (Figure 1C).

To analyse the circRNA bsj, Sanger sequencing of each gel-purified PCR amplicon was performed (Figure 1D). Given that the multiple cloning site where each of the circRNA exons were inserted is flanked by strong splice site sequences, we unsurprisingly identified distinct molecular scars at the bsj of all the mature circRNAs investigated, including residual restriction site sequence. This was confirmed in two additional cell lines – HeLa and HepG2 cells – to demonstrate this was not a cell line-specific phenomenon (Supplementary Figures S1A-B). Furthermore, the scarred bsj was not limited to an individual or pair of restriction sites in the vector multiple cloning site as it occurred when we used either the outermost or more internal pairs of restriction sites to clone the exon sequences to be circularised (Figure 1D).

When each of the three EGFR circRNAs tested above were cloned using the native intron-flanked strategy (still using a pcDNA3.1 plasmid), the circRNA bsj was free from molecular scars (Supplementary Figs S1C-D). Although this latter cloning strategy produced circRNAs with scarless bsj’s, it lacked the convenience of the universal mc vector design. We therefore set out to redesign the mc vector to ensure *bona fide* circRNAs were produced with their natural, scarless bsj at high efficiency.

### mc2, a universal vector for circRNA expression with scarless back-splice junctions

As the multiple cloning site of the mc vector was the source of the molecular scar in the circRNA bsj, we first replaced the vector’s eight restriction sites (Figures 1A-B) with a pair of inverted *Bsm*BI restriction sites (Figures 2A-B). These were oriented and positioned such that restriction digestion with *Bsm*BI removed all unwanted vector DNA, leaving only the flanking *ZKSCAN1* intron sequences to facilitate circularisation (Supplementary Figure S2). Two strategies exist for inserting the circRNA exon sequences into this new vector, which is called mc2, with both amenable to using oligonucleotides or synthetic DNA fragments (ie. gBlocks® gene fragments; see Supplementary Figure S2 for details and a specific example):

**Figure 2.**
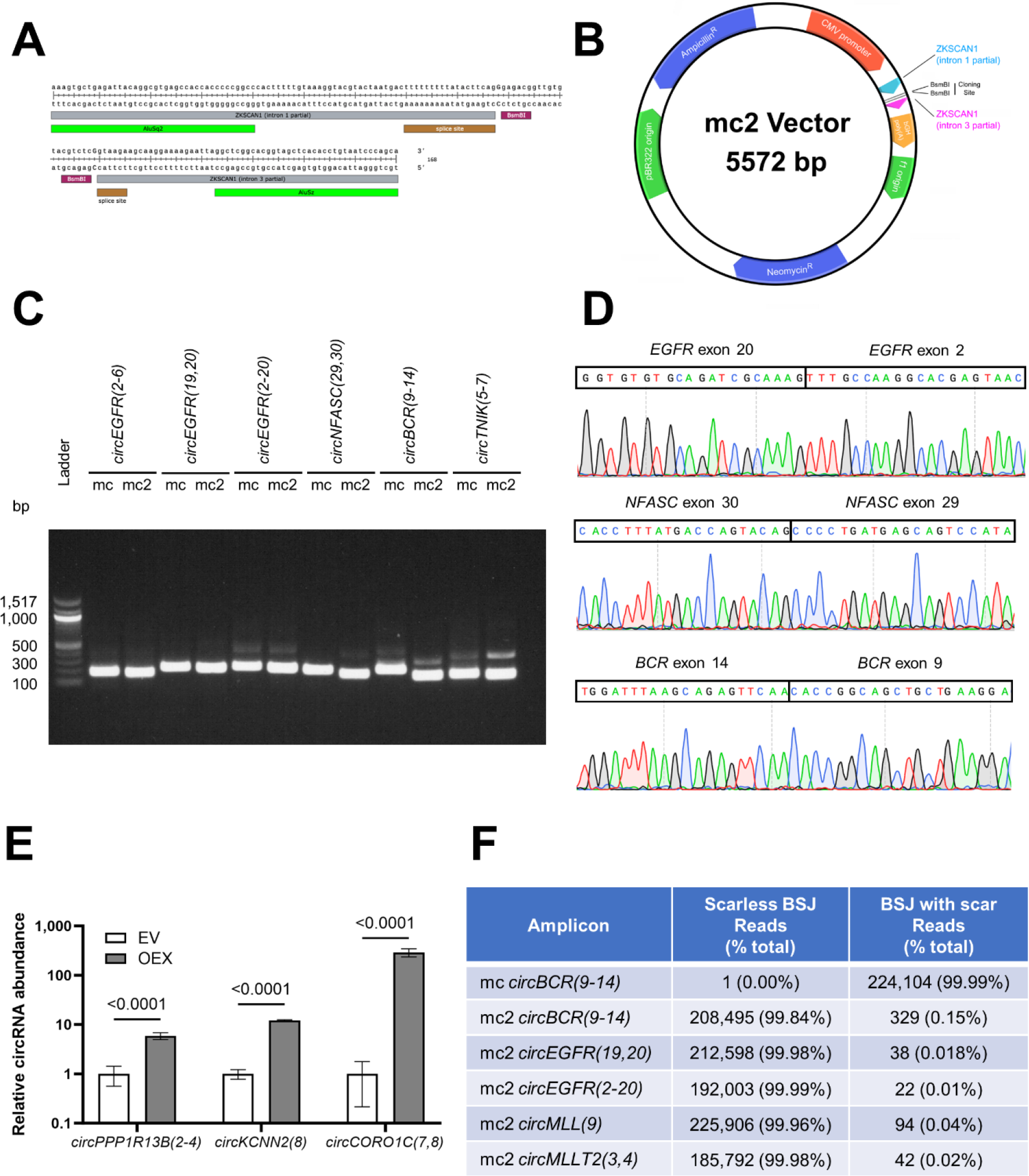
mc2, a universal vector for producing circRNAs with scarless back-splice junctions. **(A)** Nucleotide sequence of the mc2 vector including *Bsm*BI cloning sites and flanking intronic regions directing circularisation. (**B**) Plasmid map of mc2 vector generated using PlasMapper 3.0 (34). **(C)** Comparison of circRNA-specific RT-PCR products produced from HEK293 cells transfected with either mc-based or mc2-based plasmids visualised by agarose gel electrophoresis. The scarless back-splice junctions of circRNAs produced from the mc2 vector result in shorter RT-PCR amplicons. **(D)** Sanger sequence of the back-splice junction region of RT-PCR amplicons from circRNAs produced from mc2 plasmids for *circEGFR(2-20)* (top), *circNFASC(29,30)* (middle) and *circBCR(9-14)* (bottom). **(E)** qRT-PCR on RNA from cell lines following stable overexpression (OEx) of mc2-encoded circRNAs – *circKCNN2(8)* and *circPPP1R13B(2-4)* in U87 cells and *circCORO1C(7,8)* in HEK293 cells. Performed in biological and technical triplicate. nd: not detected. Data shown as mean ± SD relative to empty vector (EV). **(F)** Summary of the circRNA bsj-seq analysis of circRNAs produced by mc and mc2 plasmids. Data presented as reads for scarless (clean) and scarred bsjs and the percentage of total for each category.

1. Exons are amplified that incorporate *Bsm*BI restriction sites at the termini and are ligated into *Bsm*BI-digested mc2 vector by T4 DNA ligase.
2. Exons are amplified that incorporate 20 bp of flanking sequence for cloning by homologous recombination (ie. Gibson assembly) into *Bsm*BI-digested mc2 vector. This is particularly useful when the circRNA exons contain a *Bsm*BI site.

CircRNAs expressed from the original mc vector and the mc2 plasmid produced RT-PCR amplicons from RNase R-treated RNA. RT-PCR from HEK293 cells expressing either construct yielded a single dominant product, with the mc2-encoded circRNAs consistently producing a slightly shorter amplicon than the mc vector (Figure 2C). Sanger sequencing confirmed that, in all cases, these mc2 encoded circRNAs produced circRNA transcript with a scarless bsj (Figure 2D). A panel of 10 circRNAs were cloned into the mc2 vector and either stably, or transiently expressed in a range of human cell lines, including HEK293, HeLa, HepG2, and U87 cells confirming the bsj as scarless in each instance (Supplementary Figure S3). While 7 of these circRNAs were not endogenously expressed in the cell line we expressed them in, three of these were endogenously expressed with qRT-PCR demonstrating 5.8-290-fold overexpression in these lines (Figure 2E).

To more thoroughly investigate the fidelity of the circRNA bsj sequence produced from mc2 in cells transfected with this plasmid, a next generation sequencing strategy was devised for each individual circRNA, called circRNA bsj-seq. This was performed on five different circRNAs, including *circBCR(9-14)* cloned into both mc2 and the original mc vector in addition to another four circRNAs: *circEGFR(19,20), circEGFR(2-20), circMLL(9)* and *circMLLT2(3,4)*. Using a 12-mer unique molecular identifier, at least 185,000 individual circRNAs were sequenced from HEK293 cells transfected with each different mc/mc2 construct. It was found that only one read of *circBCR(9-14)* expressed from the mc vector was produced with a scarless bsj, while over 220,000 unique circRNA transcripts possessed a molecular scar including restriction enzyme sites. In contrast, *circBCR(9-14)* cloned into mc2 produced over 208,000 scarless bsj reads, with 0.15% possessing a molecular scar (Figure 2F).

The trend was consistent for all four other circRNAs cloned into mc2, confirming that the cloning site refinements of mc2 produce circRNAs with high fidelity back-splice junctions. This is especially important as HEK293 cells naturally express *circMLLT2(3,4)* but not the other circRNAs listed (21) and despite the circRNA bsj-seq methodology not being able to distinguish between endogenous and plasmid-derived circRNAs, the proportions of scarless bsj were consistently above 99.8% for all mc2-derived circRNAs (Figure 2F).

### Detrimental effect of RNA binding protein consensus sequences on mc2 circularisation efficiency

While complementary RNA sequence within introns can facilitate circRNA formation, the strategic use of RNA-binding protein (RBP) sites can increase circularisation and, in some cases, stimulate the biogenesis of unique circRNAs (20, 24). Notable RBPs which have been identified as circRNA biogenesis factors include Quaking (QKI) (20), fused in sarcoma (FUS) (26), Nova-2 (27) and muscleblind-like splicing regulator 1 (MBNL1) (28). To investigate what effect the presence of binding sites for these factors might have on the efficiency of circRNA formation by mc2, the *Bsm*BI sites were flanked with direct repeats of binding sites for each of these RNA-binding proteins, individually and with all four in combination (*Poly*, Figure 3A). These plasmids were Sanger sequence-verified and the resultant mc2 circRNA constructs transfected into HEK293 cells.

**Figure 3.**
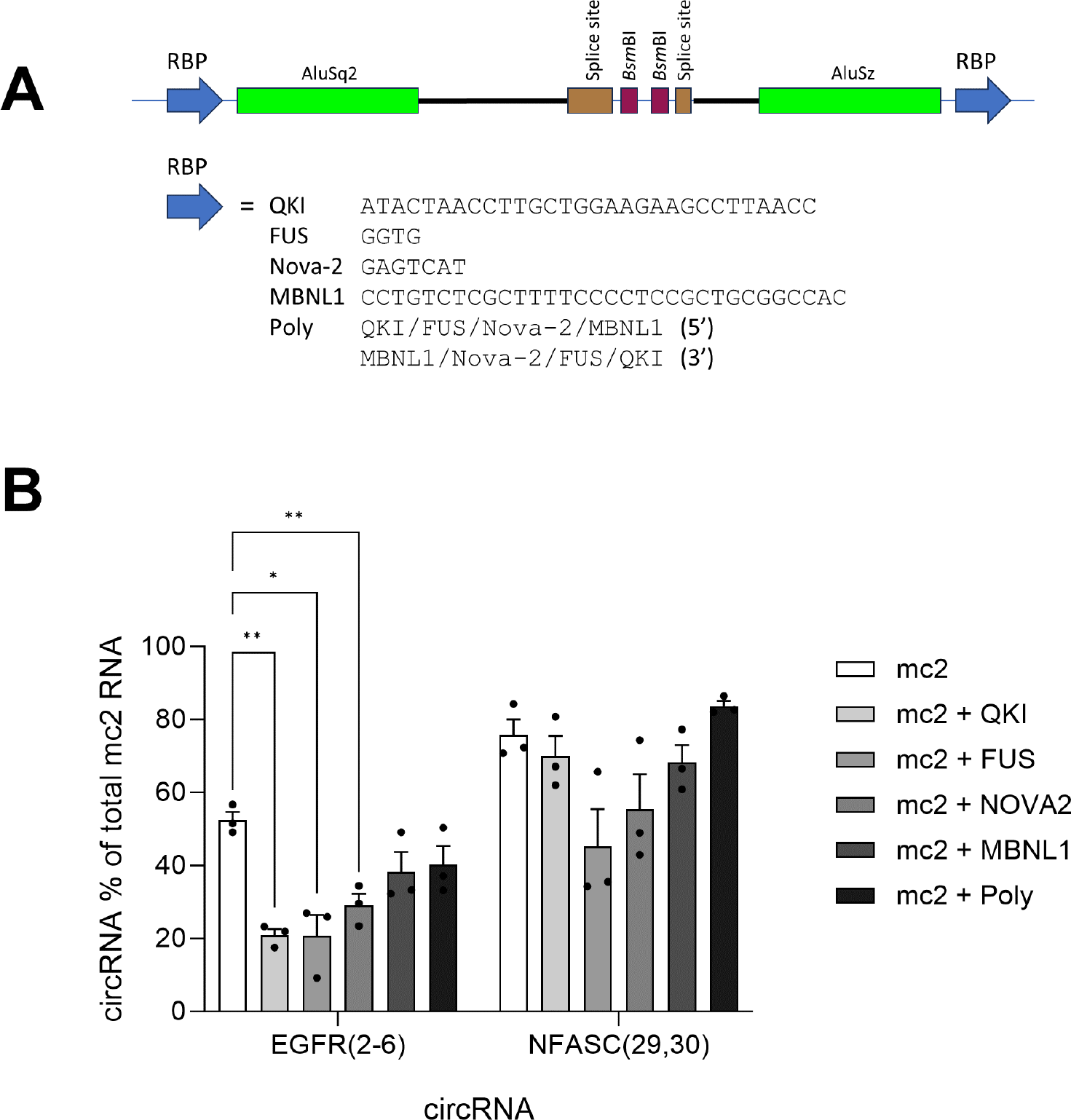
mc2 can produce circRNA with high efficiency which is not increased with the addition of RNA binding protein binding sites. (**A**) Design of the mc2 plasmid variants containing Quaking (QKI), fused in sarcoma (FUS), Nova-2, and muscleblind-like splicing regulator 1 (MBNL1) RNA binding protein (RBP) binding site sequences. (**B**) Steady state circularisation efficiency of circRNAs produced from mc2 with each of the RNA binding protein binding sites is shown compared to mc2 neo. Data is presented as circularisation efficiency established by qRT-PCR of the amount of circRNA measured, expressed as a percentage of the sum of circRNA and linear RNA. Bars show the mean±standard error of the mean for three technical replicates of single RT-qPCR experiments. Significance was determined by two-way ANOVA using Dunnett’s multiple comparisons test. Only significant differences are shown, * <0.05, ** <0.01.

The circularisation efficiency (CE) of two circRNAs – *circEGFR(2-6)* and *circNFASC(29,30)* - across each of these plasmids was calculated as the relative abundance of circular RNA transcript (qRT-PCR using circRNA specific, divergent primers) against total linear RNA transcribed from mc2 (qRT-PCR using linear exon-specific forward oligonucleotide and vector-specific reverse oligonucleotide) in a single cell line. The parent mc2 plasmid exhibited one of the highest steady-state circularisation efficiencies reported (without the use of transcriptional blockers like actinomycin D) with *circEGFR(2-6)* found to have a CE of 52%, while *circNFASC(29-30)* had a CE of 74% (Figure 3B). Interestingly, despite three of the four RNA binding proteins being expressed abundantly by HEK293 cells (Supplementary Figure S4), we found the addition of cognate RBP binding sites to the mc2 vector did not enhance the efficiency of circRNA expression for both circRNAs and, in some instances, caused a statistically significant reduction (Figure 3B). Thus, the circularisation efficiency of mc2 was not improved by the addition of RNA binding sites for FUS, QKI, Nova-2 and/or MBNL1.

### Functional validation of mc2-encoded circRNAs

To assess whether the circRNAs produced from the mc2 plasmid are functional, a luciferase reporter assay was developed which exploited the capacity of specific circRNAs to base pair with DNA and form R-loops (circR-loop) and inhibit transcription at the precise site of base-pairing (21, 29, 30). Mining our previous data in HEK293 cells where DNA:RNA immunoprecipitation was performed using the S9.6 antibody, it is clear that *circMLLT2(3,4)*, forms a strong R-loop at its cognate locus (exons 3 – 4) (Figure 4A) (21). Thirty nucleotides spanning the bsj were inserted between the transcription start site of the SV40 early promoter and *Renilla* luciferase of the psiCheck2 plasmid (Figure 4B). Acting as an internal control, firefly luciferase is encoded on the same plasmid under the control of the HSV-TK promoter, permitting quantitation of the transcriptional impact on *Renilla* luciferase driven by the circR-loop. For *circMLLT2(3,4)*, both the scarred bsj (containing *Sac*II and *Eco*RV sites, Supplementary Figure S3) and scarless bsj (Supplementary Figure S3) were cloned into separate psiCHECK2 vectors and HEK293 cells co-transfected with mc or mc2-expressing *circMLLT2(3,4)*. Comparing the normalised *Renilla*:firefly luciferase ratio of mc2-*circMLLT2(3,4)* to the control there was a 71% decrease in luciferase assay signal, confirming the transgenic circRNAs encoded by mc2 are functional. Similarly, mc-encoded *circMLLT2(3,4)* saw a significant 32.4% decrease in luciferase signal using the scarred bsj in psiCHECK2 compared with control (Figure 4C; one-way ANOVA, P = 0.0027).

**Figure 4.**
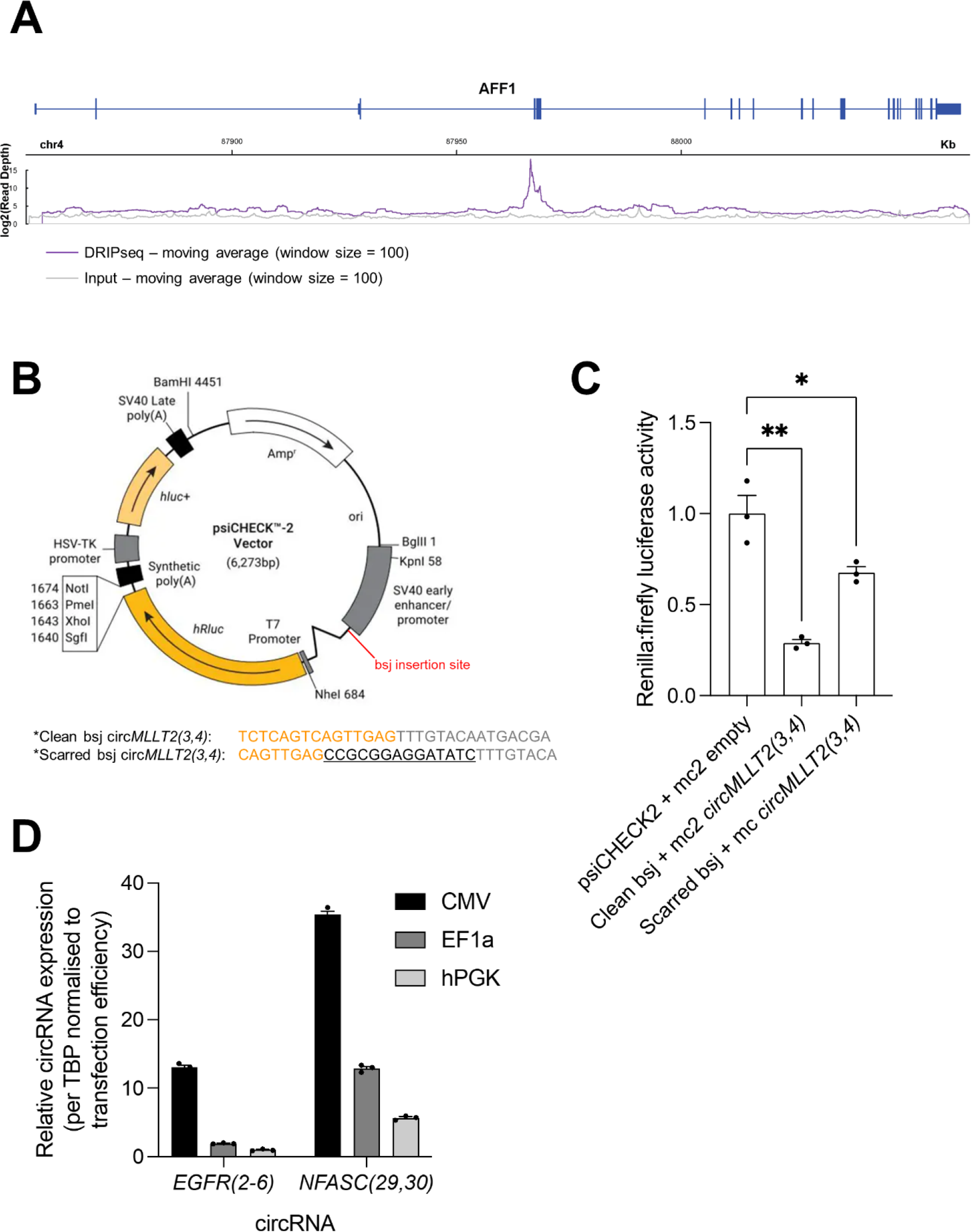
mc2-encoded circRNAs are functional. (**A**) DRIP-seq trace (purple) in HEK293 cells compared with input (grey) for the *AFF1* (*MLLT2*) gene showing a clear peak over exons 3-4 where *circMLLT2(3,4)* is bound. (**B**) Dual luciferase assay reporter vector with clean and scarred bsj sequences for *circMLLT2(3,4)* cloned between the SV40 promoter and *Renilla* luciferase, immediately downstream of the *Stu*I restriction site (shown in red). Bsj sequences consist of *MLLT2* exon 4 sequence (orange), *MLLT2* exon 3 sequence (grey), and residual mc vector restriction site sequence (black, underscored). (**C**) Luciferase reporter assay result for co-transfection of luciferase reporter with (i) control: psiCHECK2 with scarred bsj with empty mc vector, (ii) psiCHECK2 with clean bsj + mc2-circMLLT2(3,4) and (iii) psiCHECK2 with scarred bsj and mc-circMLLT2(3,4) plasmids into HEK293 cells. Data presented as mean ± SEM for three replicates, normalised to psiCHECK2 control. Significance was determined by two-way ANOVA using Dunnett’s multiple comparisons test. Only significant differences are shown, * <0.05, ** <0.001. (**D**) Variants of mc2 for expressing different levels of circRNA. HEK293 cells were transfected with mc2 plasmids expressing two different circRNAs - *circEGFR(2-6)* and *circNFASC(29,30)* - from either the human cytomegalovirus (CMV) immediate early enhancer/promoter, human phosphoglycerate kinase 1 (hPGK) promoter, or the human elongation factor 1a (EF1a) promoter. RT-qPCR was performed using RNA harvested 24 hours later. CircRNA expression is shown relative to TATA binding protein (*TBP*) expression, normalised to transfection efficiency using plasmid-encoded β-lactamase expression. Bars show the mean±standard error of the mean for three technical replicates of single RT-qPCR experiments.

### A universal toolkit for expressing circRNAs with their native back-splice junction

The circRNA vector mc2 is a derivative of pcDNA3.1, a plasmid which confers resistance to the antibiotic G418 via the *Neo*^*R*^ gene. While this is not important for transient expression of circRNAs, it enables cells transfected with this plasmid to be selected from non-transfected cells for longer term investigation of circRNA expression. To provide alternative ways to select for stably transfected cells, and thereby increase the utility of mc2, we substituted the *Neo*^*R*^ gene with genes that allow selection of transfected cells using puromycin, hygromycin B or blasticidin S and red (mCherry) or green (mNeon) fluorescence (Supplementary Figure S5). Additionally, we substituted the strong cytomegalovirus (CMV) enhancer/promoter of pcDNA3.1 with either the weaker elongation factor 1 alpha (EF1a) core promoter or the human phosphoglycerate kinase 1 (hPGK) promoter for less supraphysiological expression of circRNAs (Figure 4D) and/or expression of circRNAs in cells where the CMV enhancer/promoter is not functional. Furthermore, as some cells are difficult to transfect, we created a set of lentiviral plasmid-based variants (using the pF CAG luc puro plasmid – AddGene #67501 (31)) of mc2 to permit lentivirus-mediated transduction to achieve circRNA overexpression. All plasmids were validated by Oxford nanopore long range sequencing and have been deposited with Addgene for distribution (Table 1).

**Table 1.**
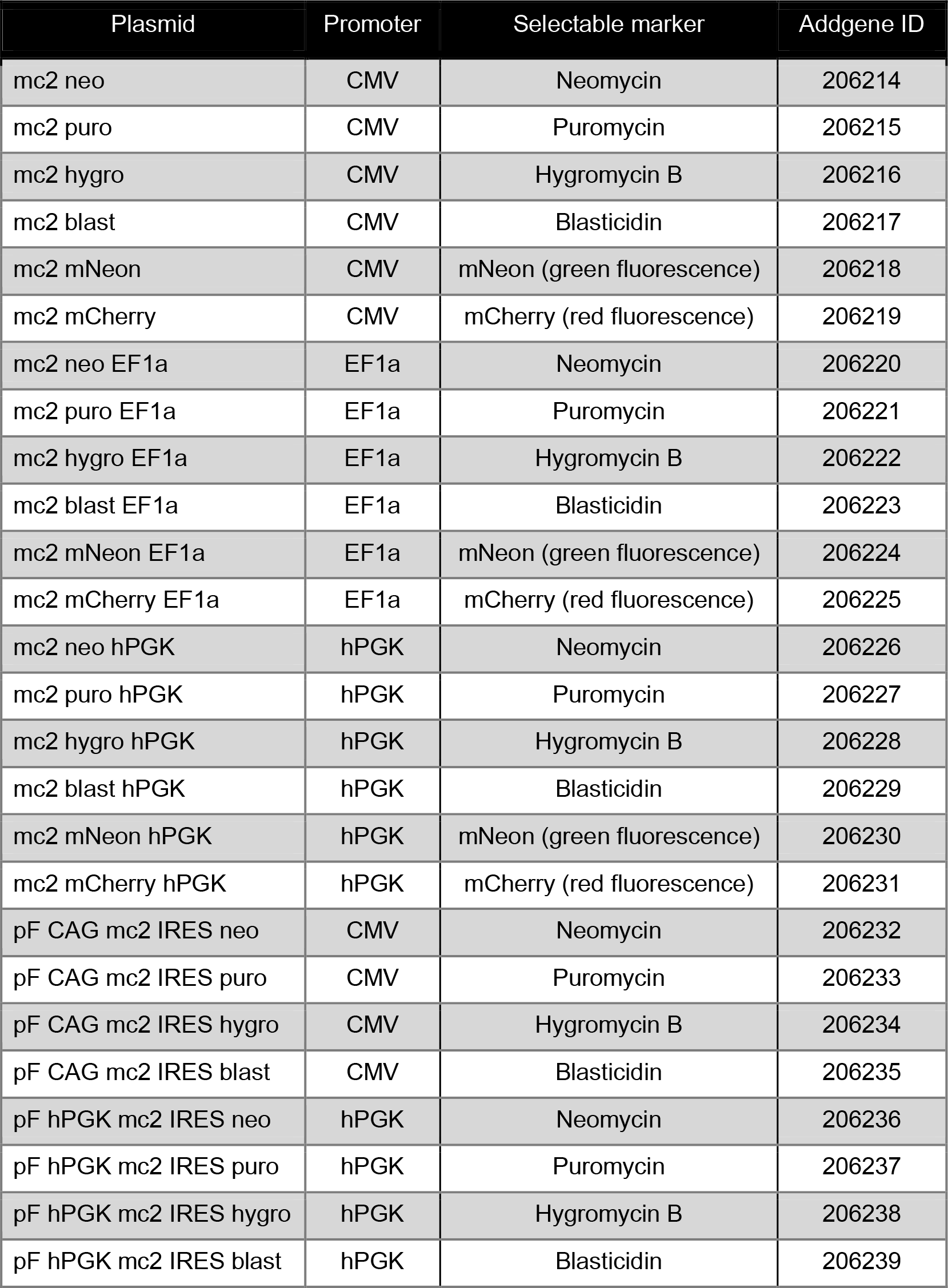
mc2 plasmids available from AddGene. Promoters are cytomegalovirus (CMV) enhancer/promoter, elongation factor 1 alpha (EF1a) core promoter or the human phosphoglycerate kinase 1 (hPGK) promoter.

## DISCUSSION

A critical approach in the investigation of circRNA function is to overexpress them in a chosen cellular context and evaluate the consequences. A few methods are routinely employed to achieve this, yet those in common use can be time-consuming, not amenable to high throughput cloning strategies or, as outlined here, be fraught with potentially functionally detrimental molecular scars. This re-engineered plasmid, mc2, corrects each of these issues. The usefulness of this strategy is extended by providing a versatile toolkit with a range of promoters, selectable markers and lentiviral options for high-efficiency and scarless expression of circRNAs for the investigation of circRNA function. While we concede that circRNA function is not uniquely limited to the fidelity of the bsj sequence, there are clear examples, including the incorporation of microexon sequences into the bsj and body of the circRNA (32) and the existence of open reading frames which span the bsj (16), where the integrity of the bsj can affect the complexity, structure and potentially function of numerous circRNAs. For these reasons, validating the bsj of the circRNA transcript along with the restriction sites used to clone and oligonucleotides used to amplify and detect these circRNAs should be standard reporting for circRNA research (33). For these reasons, we encourage the circRNA research field to adopt these mc2 plasmids for greater confidence in experiments involving transgenic overexpression of scarless circRNAs.

## DATA AVAILABILITY

The mc2 plasmids described are available from Addgene (https://addgene.com, #206214-206223). CircRNA back-splice junction amplicon-seq data has been deposited with the GEO repository with accession number GSE246020 (reviewer token: sboluoqcfnopjkt). DRIP-seq data has been deposited with the GEO repository with accession number GSE125985.

## SUPPLEMENTARY DATA

Supplementary Data are available at NAR online.

## ACKNOWLEDGEMENT

The authors gratefully acknowledge the South Australian Genomics Centre (SAGC) for amplicon-seq library preparation and sequencing. The SAGC is supported by the National Collaborative Research Infrastructure Strategy (NCRIS) via BioPlatforms Australia and by the SAGC partner institutes.

