## Supplementary Figures for "Versatile toolkit for highly-efficient and scarless overexpression of circular RNAs"

### Supplementary Figure S1

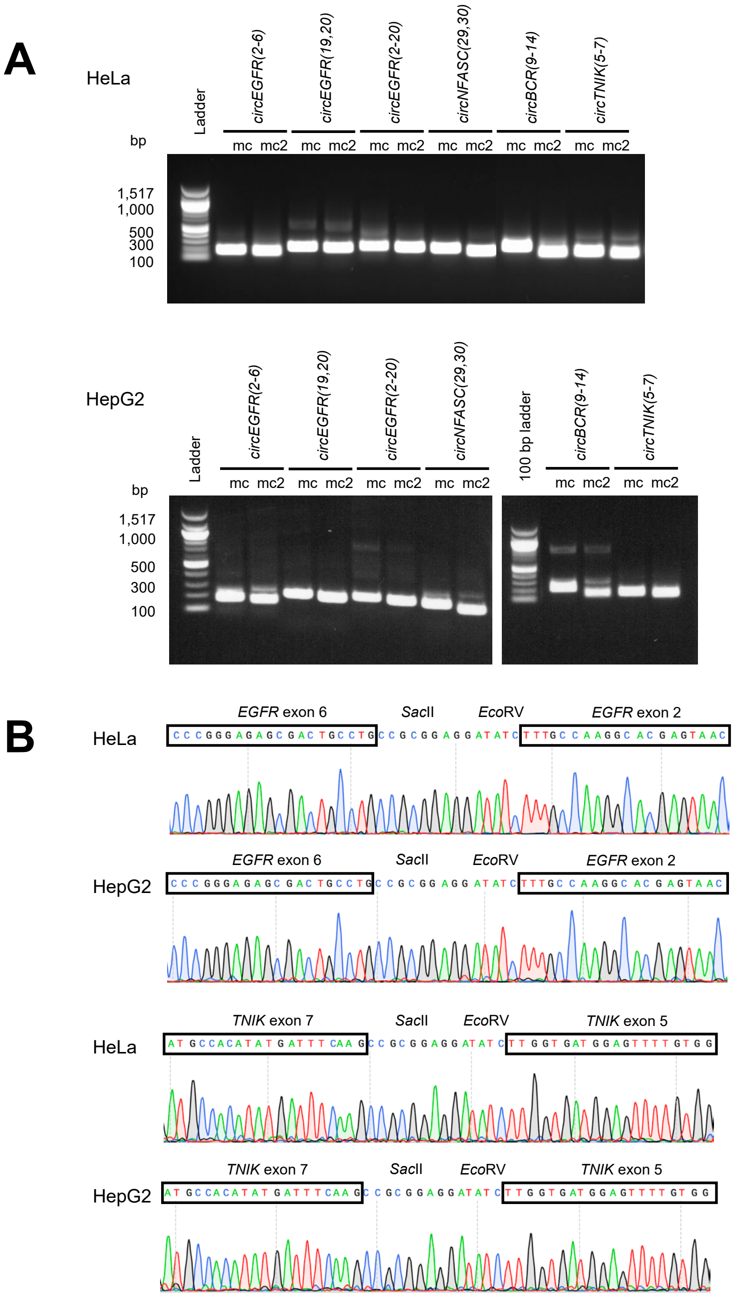


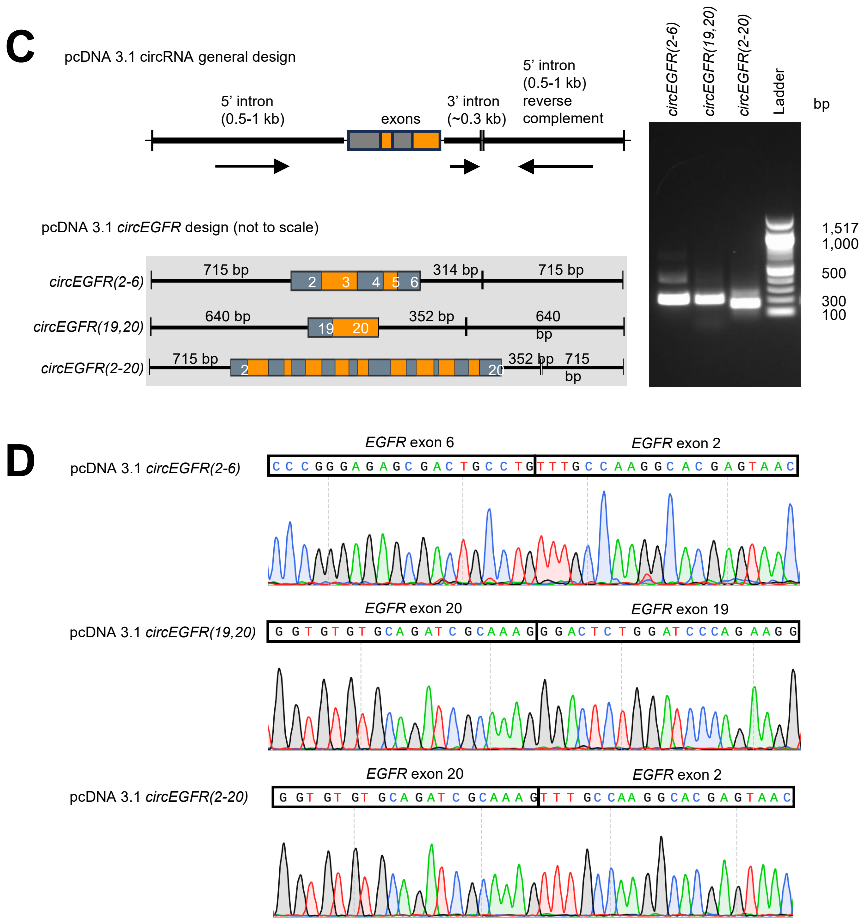


**Supplementary Figure S1.** CircRNA Mini Vector (mc) incorporates vector multiple cloning site sequence within the back-splice junction of expressed circRNAs but alternative vector designs do not. **(A)** Comparison of circRNA-specific RT-PCR products produced from HeLa (top) and HepG2 (bottom) cells transfected with either mc- or mc2-based plasmids, visualised by agarose gel electrophoresis and SYBR Safe DNA gel stain. RNA was isolated 24 hours post transfection. The larger RT-PCR amplicons produced from mc vector plasmids contain unwanted vector sequences from the multiple cloning site within the back-splice junctions of the circRNAs produced. **(B)** Sanger sequence of the back-splice junction region of RT-PCR amplicons from HeLa and HepG2 cells transfected 24 hours previously with mc vector clones of *EGFR* exon 2-6 (top) and *TNIK* exons 5-7 (bottom). **(C)** An alternative circRNA vector design incorporating inverted native intron sequences successfully produced circRNAs with scarless back-splice junctions. The left panel shows the general vector design of pcDNA 3.1 circRNA (top) and the specific arrangement of three constructs for expressing circRNAs from the *EGFR* gene. The right panel shows circRNA-specific RT-PCR products from HEK293 cells transiently transfected with the three circRNA-expressing *EGFR* plasmids. **(D)** Sanger sequence of the back-splice junction region of the RT-PCR amplicons from HEK293 cells transfected 24 hours previously with the pcDNA 3.1 circRNA clones of *EGFR* exon 2-6 (top), *EGFR* 19-20 (middle), and *EGFR* exons 2-20 (bottom).

### Supplementary Figure S2

#### mc2 cloning strategies

mc2 vector nucleotide sequence in the region of the *Bsm*BI restriction sites:

intron sequence *Bsm*BI sites

```
…GTACTAATGACTTTTTTTTTATACT TCAGGGAGACGGTTGTGTACGTCTCG GTAAGAAGCAAGGAAAAGAATTAGGCTCG…
…CATGATTACTGAAAAAAAAATATGAAGTC CCTCTGCCAACACATGCAGAGCCATT CTTCGTTCCTTTTCTTAATCCGAGC…

…GTACTAATGACTTTTTTTTTATACT      BsmBI-digested mc2    GTAAGAAGCAAGGAAAAGAATTAGGCTCG…
…CATGATTACTGAAAAAAAAATATGAAGTC                             CTTCGTTCCTTTTCTTAATCCGAGC…
```

Four potential ways to introduce nucleotide sequence to be circularised into *Bsm*BI-digested mc2:

(1) Paired oligos (anneal oligos and ligate with *Bsm*BI-digested mc2)

5′-TCAG NNNNNNNNNNNNNNNNNNNNNNNNNNNNNNN-3′ (where N…= exon sequence to be circularised)
3′-DDDDDDDDDDDDDDDDDDDDDDDDDDDDDDD CATT-5′ (where D is complementary to N)

(2) PCR product with *Bsm*BI ends (digest with *Bsm*BI and ligate with *Bsm*BI-digested mc2)

Forward primer: 5′-NCGTCTCN TCAG $(N_{17-20})$-3′
Reverse primer: 5′-NCGTCTCN TTAC $(N_{17-20})$-3′

(3) PCR product with 20 bp of terminal sequence matching the vector ends (homologous recombination)

Forward primer: 5′-ACTTTTTTTTTATACTTCAG $(N_{17-20})$-3′
Reverse primer: 5′-TTCTTTTCCTTGCTTCTTAC $(N_{17-20})$-3′

(4) gBlock with 20 bp of terminal sequence matching the vector ends (homologous recombination)

5′-ACTTTTTTTTTATACTTCAG NNNNNNNNNNNNNNNNNNNN GTAAGAAGCAAGGAAAAGAA-3′

For example, to clone *EGFR* exon 5 (nucleotide sequence shown below) into mc2 to produce *circEGFR(5)* circRNA:

5′-GCCAAAAGTGTGATCCAAGCTGTCCCAATGGGAGCTGCTGGGGTGCAGGAGAGGAGAACTGCCAGAAAC-3′
3′-CGGTTTTCACACTAGGTTCGACAGGGTTACCCTCGACGACCCCACGTCCTCTCCTCTTGACGGTCTTTG-5′

Do any of the following:

(1) Order the following oligos (standard desalted oligos) to anneal and then ligate with *Bsm*BI-digested mc2:

Forward oligo: 5′-TCAG GCCAAAAGTGTGATCCAAGCTGTCCCAATGGGAGCTGCTGGGGTGCAGGAGAGGAGAACTGCCAGAAAC-3′
Reverse oligo: 5′-TTAC GTTTCTGGCAGTTCTCCTCTCCTGCACCCCAGCAGCTCCCATTGGGACAGCTTGGATCACACTTTTGGC-3′

(2) Order the following oligos to amplify the exon by PCR from gDNA or cDNA, digest it with *Bsm*BI, and ligate it with *Bsm*BI-digested mc2 :

Forward primer: 5′-GCGTCTCA TCAG GCCAAAAGTGTGATCCAAGC-3′
Reverse primer: 5′-TCGTCTCA TTAC GTTTCTGGCAGTTCTCCTCT-3′

(3) Order the following oligos to amplify the exon by PCR from gDNA or cDNA, then clone it into *Bsm*BI-digested mc2 using an NEBuilder kit:

Forward primer: 5′-ACTTTTTTTTTATACTTCAG GCCAAAAGTGTGATCCAAGC-3′
Reverse primer: 5′-TTCTTTTCCTTGCTTCTTAC GTTTCTGGCAGTTCTCCTCT-3′

(4) Order the gBlock shown below and clone it into *Bsm*BI-digested mc2 using an NEBuilder kit:

5′- ACTTTTTTTTTATACTTCAG
GCCAAAAGTGTGATCCAAGCTGTCCCAATGGGAGCTGCTGGGGTGCAGGAGAGGAGAACTGCCAGAAAC GTAAGAAGCAAGGAAAAGAA-3′

**Notes for cloning into mc2**

We use any of the four methods for cloning exon sequences into the mc2 vectors to be transcribed into RNA and circularised.
Each method begins by digesting mc2 with *BsmB*I restriction endonuclease at 50°C (and then optionally dephosphorylating it with Antarctic alkaline phosphatase).
We typically digest 1 µg of mc2 for 1 hour.
While the *BsmB*I-digested mc2 plasmid can be separated from the 22 bp *BsmB*I restriction fragment by agarose gel electrophoresis and then recovered from an agarose gel slice, we routinely just recover it from the digest reaction using a DNA purification kit (e.g. QIAquick), eluting it in 30 µL of water.
DNA sequence to be transcribed into RNA, and then circularised, is cloned between the *BsmB*I ends of mc2.

(1) Very short sequences can be cloned as doubled-stranded oligonucleotides of the form shown on the preceding page.
If the mc2 plasmid was dephosphorylated, the double-stranded oligonucleotides should be phosphorylated using T4 DNA kinase.
We'd normally not dephosphorylate *Bsm*BI-digested mc2 in this case and use unphosphorylated annealed oligos instead.
*BsmB*I-digested mc2 and doubled-stranded oligonucleotides are ligated using T4 DNA ligase.

(2) Longer exon sequences, or multiple exon sequences, can be amplified from cDNA using a high-fidelity DNA polymerase (e.g. NEB Q5 DNA polymerase) and oligonucleotide primer pairs incorporating a 5' *BsmB*I restriction site (of the form shown on the preceding page).
The PCR amplicon is then digested with *BsmB*I, recovered using a DNA purification kit, and ligated with *BsmB*I-digested mc2 using T4 DNA ligase.
Check that the exon sequences do not contain an endogenous *BsmB*I restriction site if this method is used.

(3) Exon sequences also can be amplified from cDNA using a high-fidelity DNA polymerase (e.g. NEB Q5 DNA polymerase) and oligonucleotide primer pairs incorporating 20-nucleotide extensions that are homologous to the free ends of *BsmB*I-digested mc2.
The forward primer should be of the form: 5'-ACTTTTTTTTTATACTTCAG$(N)_{17-20}$-3'.
The reverse primer should be of the form: 5'-TTCTTTTCCTTGCTTCTTAC$(N)_{17-20}$-3'.
$(N)_{17-20}$ represents the nucleotide sequences of the primer pair that amplifies the exon(s) of interest.
The purified PCR product can be cloned into *BsmB*I-digested mc2 by homologous recombination (e.g. using an NEBuilder kit).

(4) Exon sequence can be synthesised as a gBlock (IDT) of the form 5'-ACTTTTTTTTTATACTTCAG-exon sequence-GTAAGAAGCAAGGAAAAGAA-3'
The gBlock can be cloned into *BsmB*I-digested mc2 by homologous recombination (e.g. using an NEBuilder kit).

**Supplementary Figure S2.** Sequence of mc2 plasmid cloning site and general cloning considerations. Specific examples provided for cloning *circEGFR(5)*.

### Supplementary Figure S3

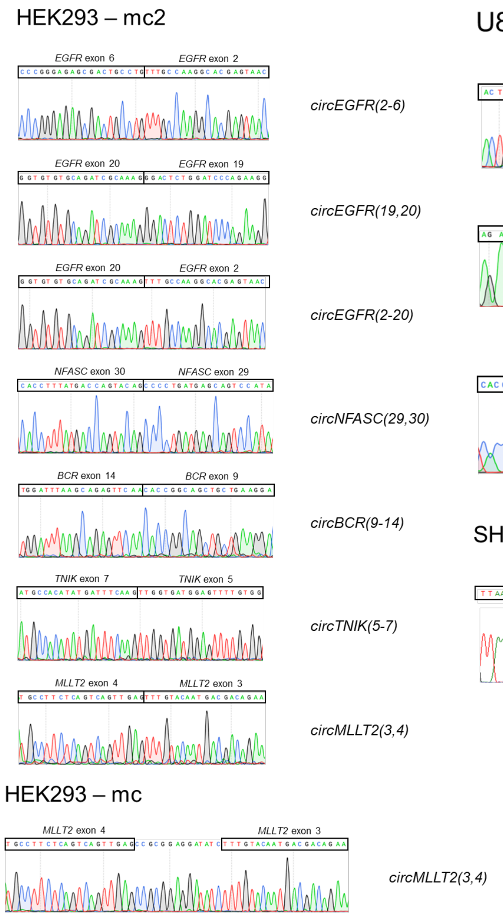


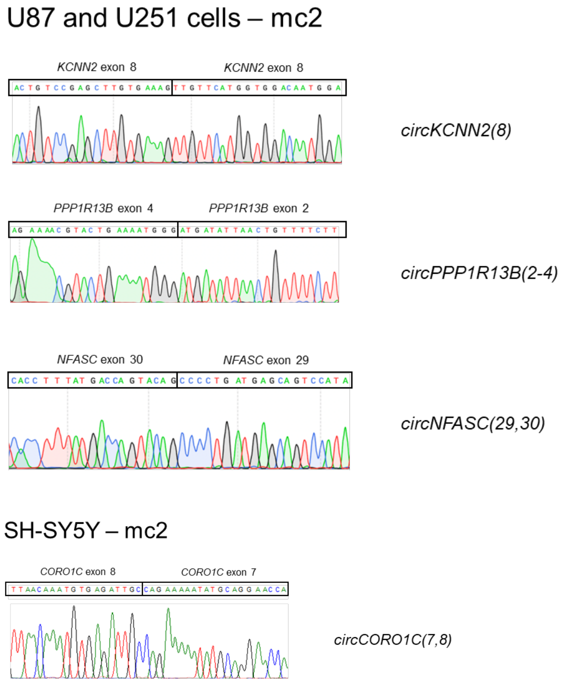


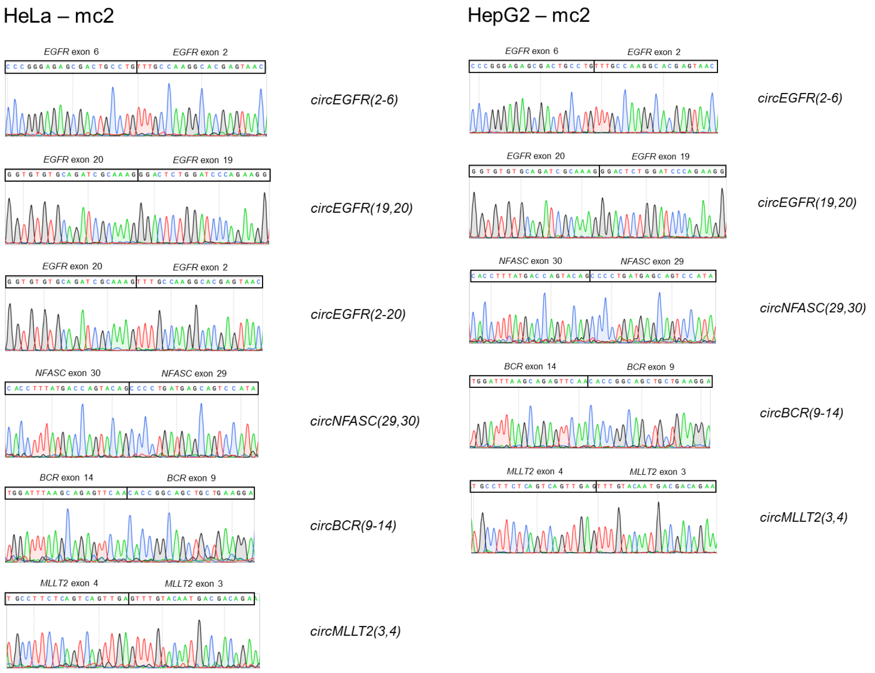


**Supplementary Figure S3.** Sanger sequence chromatograms of RT-PCR amplicons across the back-splice junction region for 20 different circRNAs from HEK293, HeLa, U87, U251, HepG2 and SH-SY5Y cells transfected 24 hours previously with mc2 vector (or mc vector).

### Supplementary Figure S4

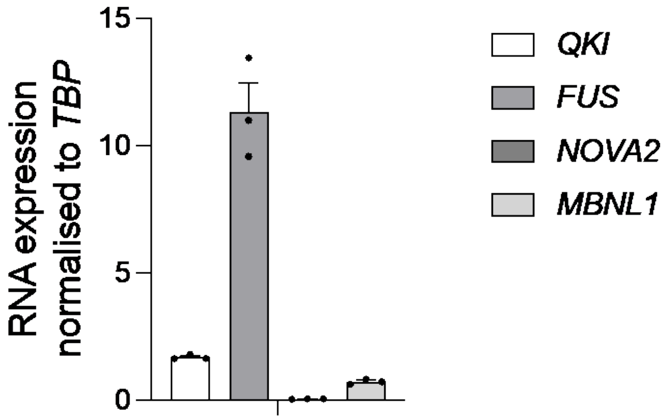


**Supplementary Figure S4.** Relative mRNA expression in HEK293 cells of the genes encoding the RNA binding proteins *Quaking* (*QKI*), *fused in sarcoma* (*FUS*), *NOVA2*, and *muscleblind-like splicing regulator 1* (*MBNL1*). qRT-PCR using gene-specific primers shown in Supplementary Table 1, mRNA expression is shown normalised to TATA binding protein (*TBP*). Data presented as mean ± standard deviation, n = 3 biological replicates.

### Supplementary Figure S5

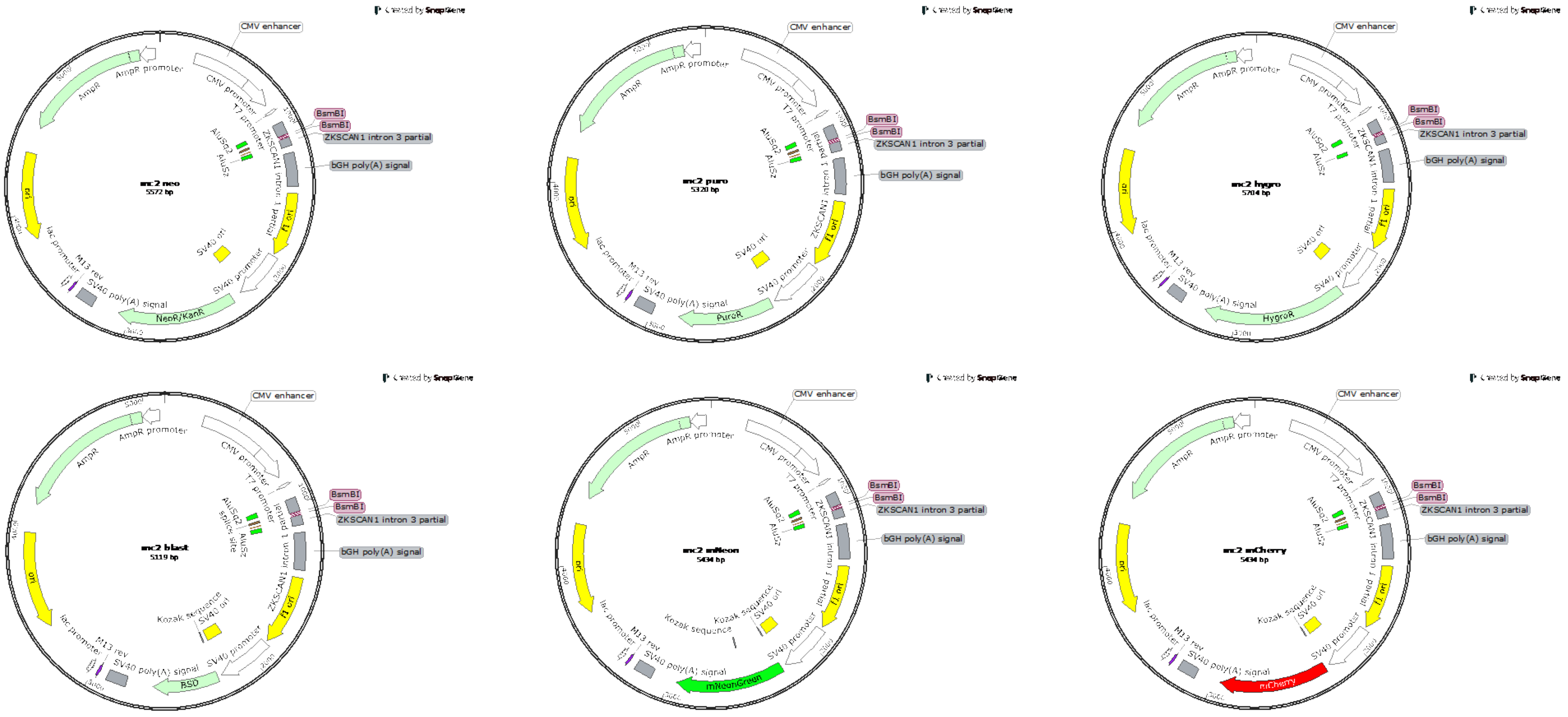


**Supplementary Figure S5.** Plasmid maps for the six human cytomegalovirus enhancer/promoter variants of mc2 for selecting stably-transfected cells using either G418 (neoR), puromycin (puroR), hygromycin B (hygroR), or blasticidin S (BSD), or green (mNeon) or red (mCherry) fluorescence.
